# Proteostasis Signatures in Human Diseases

**DOI:** 10.1101/2024.12.23.630032

**Authors:** Christine M Lim, Michele Vendruscolo

## Abstract

The protein homeostasis (proteostasis) system maintains the proteome in a healthy state. Although this system has been comprehensively mapped, its perturbations in disease remain incompletely characterised. To address this problem, here we define the proteostasis signatures, which represent the characteristic patterns of change in the proteostasis system associated with disease. We performed a large-scale, pan-disease analysis across 32 human diseases spanning 7 disease types. We first identified unique proteostasis perturbations in specific disease states. We then uncovered distinctive signatures differentiating disease types, pointing to a range of proteostasis mechanisms in disease development. Next, we tracked the temporal evolution of proteostasis signatures, revealing shifts in proteostasis disruption over the course of disease progression. Finally, we demonstrated how smoking, a major risk factor of many diseases, impairs proteostasis in a manner similar to disease, potentially creating a predisposed environment for disease onset. These results illustrate the opportunities offered by the study of human diseases from the perspective of proteostasis signatures.

---

The proteostasis system coordinates the synthesis, folding, trafficking, and degradation of proteins (1-3). This intricate network encompasses molecular chaperones, degradation pathways, and regulatory systems that collectively ensure the proper maintenance of the proteome (1-3). Maintaining the balance between these processes is essential for cellular function and organismal health, and its disruption has been implicated in numerous diseases, including cancer, neurodegenerative disorders, and autoimmune conditions (4-7). As proteins misfold, are damaged and fail to be degraded, cells face increased stress and dysfunction, contributing to disease pathogenesis (4-7). Understanding the mechanisms underlying proteostasis impairment is critical for developing therapeutic strategies that restore cellular balance (4-7).

Proteostasis dysregulation has been shown to manifest in distinct patterns across different diseases, reflecting the diversity of underlying mechanisms (1-5). For example, neurodegenerative conditions such as Alzheimer’s and Parkinson’s disease are characterized by the progressive aggregation of misfolded proteins, whereas cancers exploit proteostasis systems like the ubiquitin-proteasome pathway to sustain rapid cell division (1-5). While these disease-specific patterns are well-recognized, their broader significance as systematic signatures of proteostasis dysfunction has yet to be fully elucidated.

Inspired by the impact of the study of mutational signatures in cancer research (8-10), which have identified the molecular drivers of tumor biology and guided targeted therapies, we investigated a similar approach for understanding proteostasis disruption across diseases. Proteostasis pathways are known to be intricately linked to many diseases, each characterized by unique patterns of cellular damage (1-5, 11, 12). To capture these distinct molecular alterations systematically, we descry be the concept of proteostasis signatures, which provides a framework for linking specific proteostasis pathway disruptions to disease mechanisms.

By characterising proteostasis signatures, we mapped the proteostasis dysregulation across diseases. By defining these signatures, we aim to provide a systematic framework for understanding how proteostasis is disrupted in different disease contexts and stages. This framework has the potential to bridge gaps in our knowledge by linking specific proteostasis pathways to their functional consequences in health and disease.

## Results

### Proteostasis proteins are closely associated with disease

We first asked whether the proteins involved in proteostasis are preferentially associated with disease. To this end, we analysed a recent comprehensive map of the human proteostasis network (13, 14) and used the proteins within the network as our set of proteostasis proteins. We computed their association to 32 diseases from 7 disease groups **(Table S1)** by studying the prevalence of proteostasis proteins within the gene set of each disease (disease gene set). The method for generating each disease gene set is described in **Materials & Methods**. Our results show that proteostasis proteins are closely associated with disease, as they are significantly over-represented in disease protein sets **(Figure 1)**. Over-representation analysis of each of the 4 protein groups within the top 500 disease-associated proteins for every disease was computed with the hypergeometric test.

**Figure 1.**
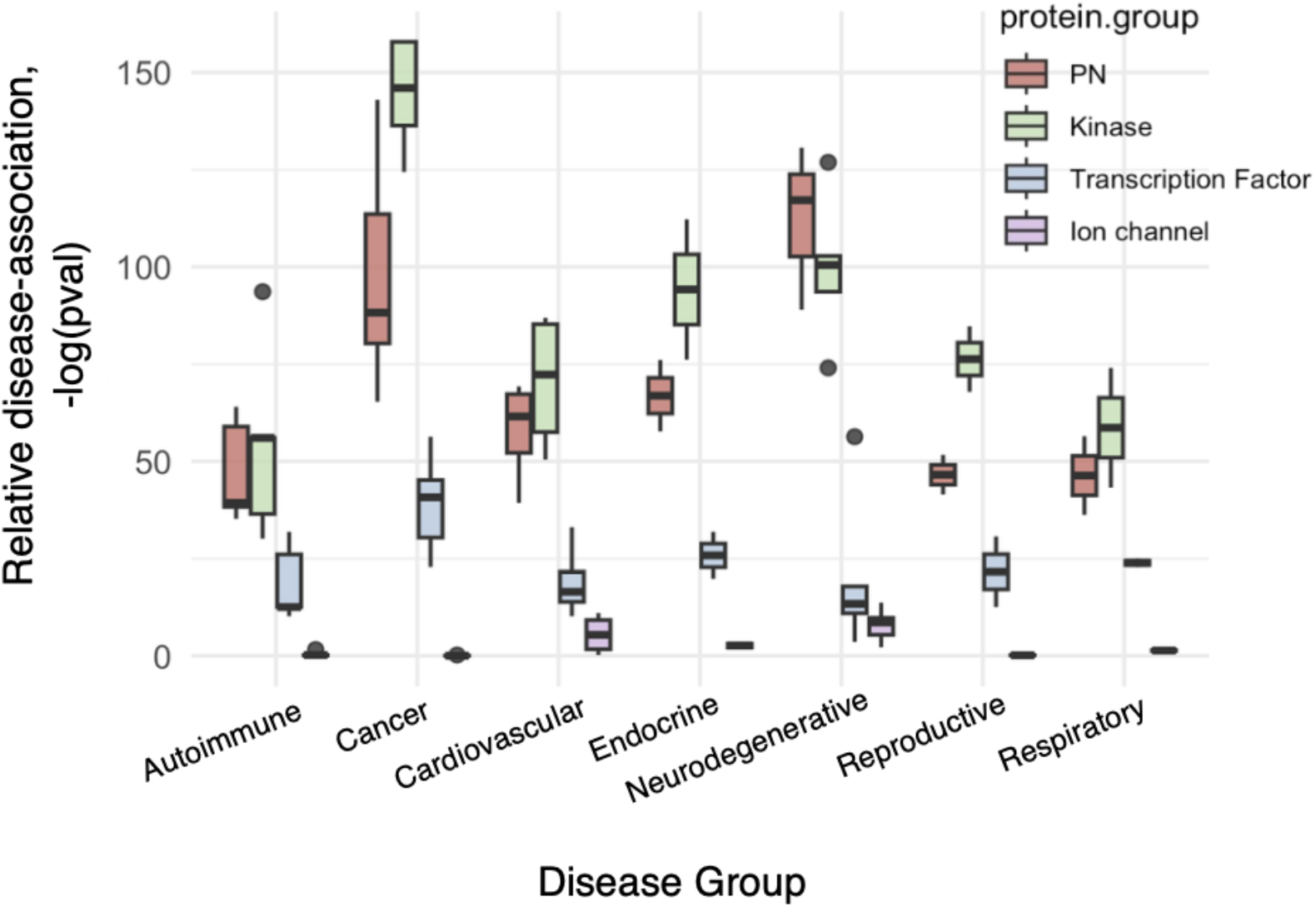
Proteostasis proteins are closely associated with disease. Relative disease association of proteostasis proteins (PN) were quantified and benchmarked against 3 control groups: kinases, transcription factors, and ion channels. Disease-association was determined by relative over-representation of a protein group within disease gene sets. This was done using the hypergeometric test measuring the statistical significance their prevalence within each disease gene set. P-values were plotted on a -log(p-value) scale, with higher values representing stronger significance. Based on this quantification, proteostasis proteins are significantly over-represented in all the disease groups studied. They are relatively more disease associated than transcription factors, and in some cases even than kinases.

We then compared the disease association of the proteostasis proteins against 3 well characterized disease-associated functional protein groups: kinases, transcription factors, and ion channels. Kinases (15, 16) and transcription factors (17, 18) are essential regulatory proteins controlling diverse events in cellular signalling and gene transcription. They were selected as positive control groups, as they have been widely reported to be implicated in a range of diseases (15-18). Ion channels are membrane proteins that regulate signal transduction across cell membranes (19, 20), and were selected as a negative control group, as only 2 of our 7 disease groups studied (cardiovascular and neurodegenerative) are commonly associated with ion channels (19-21). As expected, kinases and transcription factors are highly over-represented across the disease groups, while ion channels are over-represented in the neurodegenerative and cardiovascular disease groups **(Figure 1)**.

Our analysis reveals a strong relevance of proteostasis proteins in disease, with almost comparable disease-association with kinases **(Figure 1)**, which are a key targeted group of drug targets (22).

### Proteostasis profiles of disease

For each of the 32 diseases included in the study, we created a profile quantifying the proportion of proteostasis proteins within each disease gene set and associating the relevant proteostasis network branches and functional classes to the disease **(Figure 2)**. Based on our profiling, proteostasis proteins comprise a large portion of disease gene sets in cancer (25-36%) and neurodegenerative diseases (30-35%) **(Figure 2)**.

**Figure 2.**
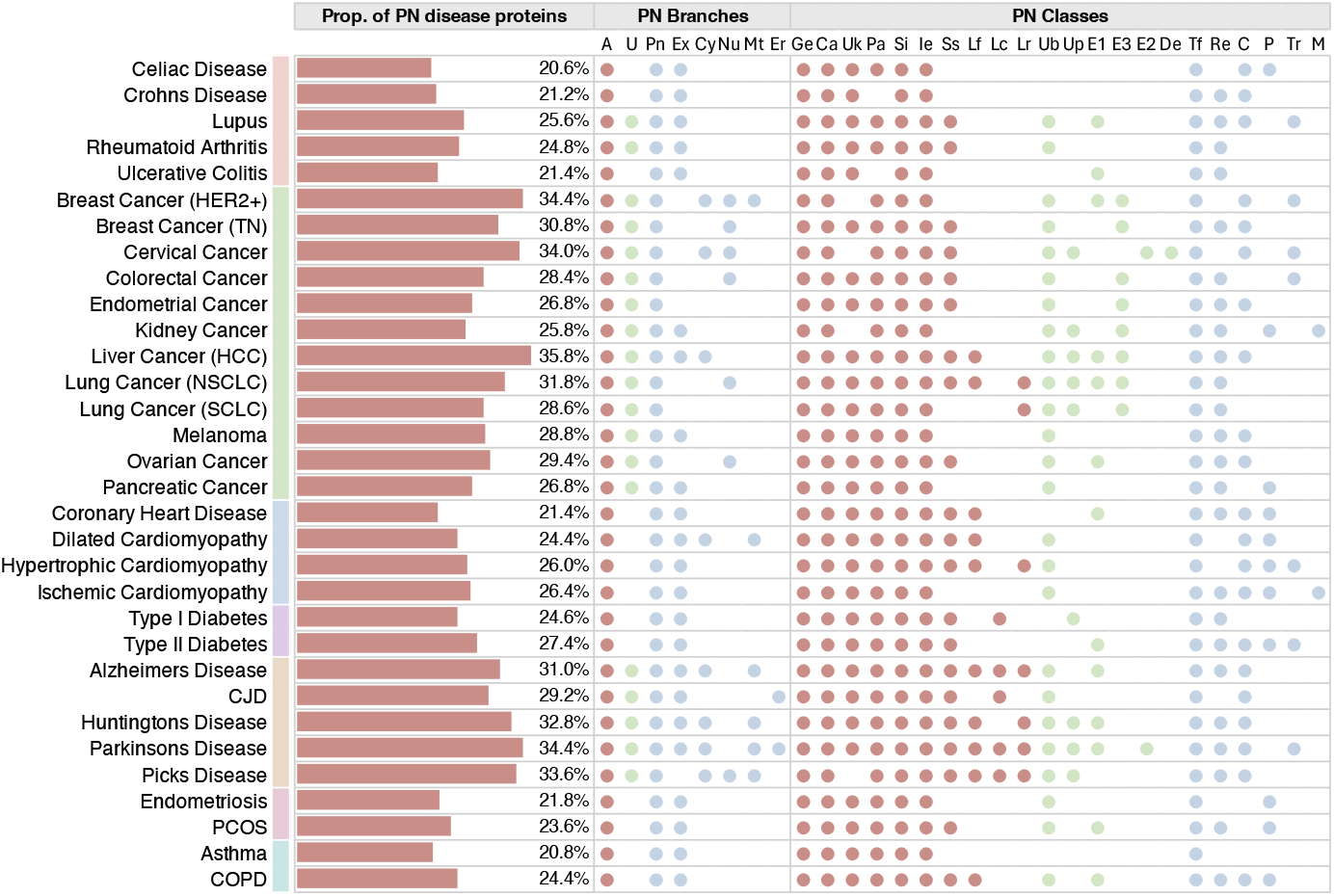
Proteostasis profiles of diseases. For each of the 32 diseases included in this study (**Table S1**, arranged in the figure by protein group), we computed the fraction of proteostasis proteins within each disease gene set (brown bars). Across the 32 diseases, the fraction of proteostasis proteins within disease gene sets ranged from 20% to 36%. We further decomposed disease genes involved in proteostasis into their relevant proteostasis branches **(Table S2)** and classes **(Table S3)**. This was done by identifying which branch/class was over-represented within the proteostais proteins corresponding to the genes of each disease gene set. The statistical significance of over-representation of a branch or class was determined using the hypergeometric test (p-value < 0.01) and represented as a spot within the figure. Colour-coding of the spots reflect their involvement in the autophagy-lysosome pathway (brown), the ubiquitin-proteasome system (green) and the anabolic system (blue).

At the branch level, proteostasis proteins involved in the autophagy-lysosome pathway (ALP) and proteostasis regulation are consistently over-represented across all diseases. We found that ubiquitin-proteasome system (UPS) proteins are closely associated with cancers and neurodegenerative diseases but not with other disease groups **(Figure 2)**. In contrast extracellular proteostasis proteins are over-represented in all disease groups except cancer **(Figure 2)**.

At the class level, most classes of the ALP machinery appear over-represented in disease gene sets, as observed at the branch level. We highlight: (a) UPS ubiquitin-binding proteins are significant in cancers and neurodegenerative diseases, while UPS E3 ligases are significantly represented only in cancers; (b) transcription factors, as reported extensively in literature, are also found to be well-represented across all disease types included in the study; and (c) molecular chaperones are a functional class with strong associations with cardiovascular and neurodegenerative diseases.

### Proteostasis states in disease

Based on the observations from the proteostasis network profiles of the diseases, we identified 3 distinct proteostasis states in disease **(Figure 3A)**. These proteostasis states describe disease in terms of characteristic perturbations of the proteostasis network. The most important branches of the proteostasis network for the definition of these states are ALP, UPS and proteostasis regulation. The first proteostasis state is characterized by significant UPS perturbation but limited involvement of extracellular proteostasis **(Figure 3A)**. This state is characteristic of cancer **(Figure 3B)**. The second proteostasis state involves extensive perturbation of both UPS and extracellular proteostasis **(Figure 3A)**. This state is largely presented in neurodegenerative diseases **(Figure 3B)**. The third proteostasis state involves the distinctive deregulation of extracellular proteostasis but limited in UPS involvement **(Figure 3A)**. This state is more common and less discriminatory, with autoimmune, endocrine, cardiovascular, reproductive, and respiratory diseases all presenting this trend.

**Figure 3.**
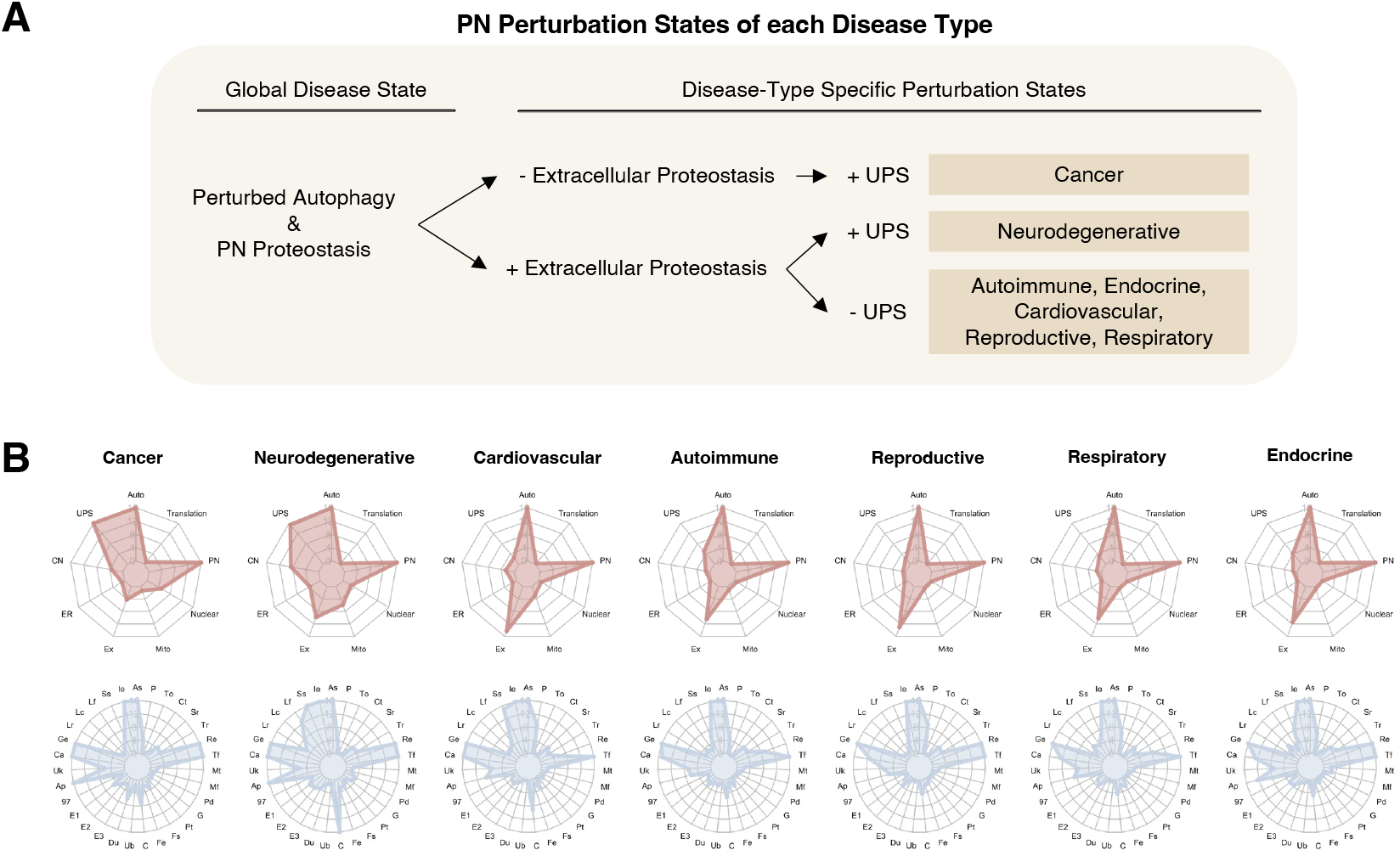
Characteristic proteostasis perturbation states and patterns in disease. **(A)** Three generalised proteostasis perturbation states capable of discriminating disease types. **(B)** Spider plot depicting trends of over-representation of the relevant proteostasis network branches and classes across all 7 disease types.

### Proteostasis signatures of disease

To further study the characteristic perturbation of the proteostasis network in a disease, we defined the proteostasis signatures of disease. We first considered 4 groups of diseases with distinct proteostasis states: cancer, neurodegenerative diseases, autoimmune diseases, and cardiovascular diseases. Clustering the diseases based on their disease-associated genes resulted in 4 clusters. The diseases clustered largely according to their proteostasis states, with a notable exception where kidney cancer and pancreatic cancer clustered with autoimmune diseases **(Figure 4A)**, a finding consistent with the bidirectional association between cancer and autoimmune disorders reported in the literature (23, 24).

**Figure 4:**
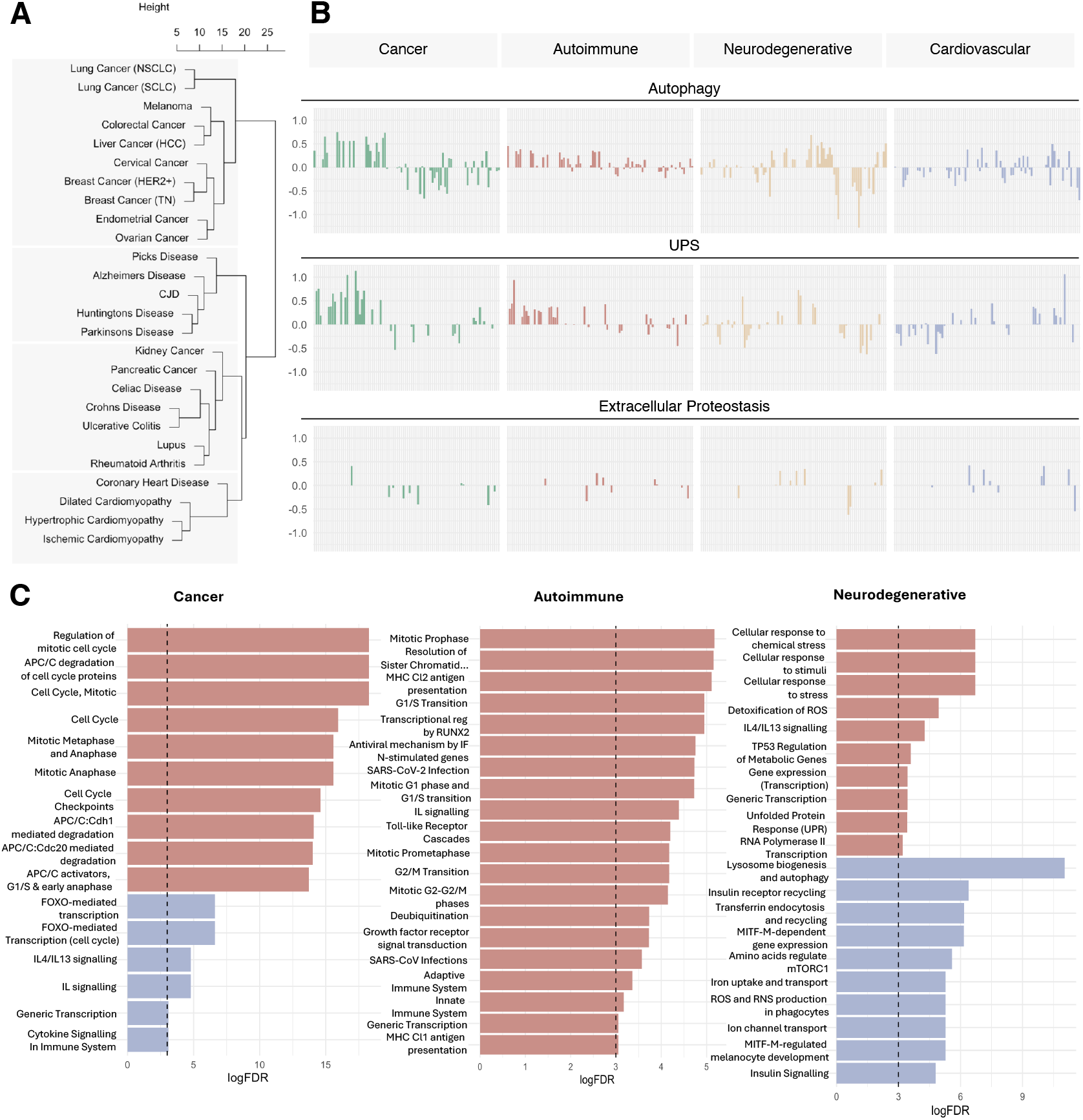
Proteostasis signatures of disease. **(A)** Unsupervised clustering of cancers, neurodegenerative diseases, autoimmune diseases, and cardiovascular disease resulted in 4 clusters. The clusters were mostly by disease type, with the notable exception of pancreatic cancer and kidney cancer clustering with autoimmune diseases. **(B)** Proteostasis signature trends reveal that cancers and autoimmune diseases have a large proportion of common genes perturbed in similar patterns. In contrast, neurodegenerative diseases are perturbed in opposite directions. **(C)** Functional implications of the proteostasis signatures. The top enriched pathways (up to 10 each) for upregulated and downregulated genes for each cluster type are shown.

Extracting generalized gene-wise disease signatures for ALP, UPS, and extracellular proteostasis identified similar proteostasis signatures ofr cancers and autoimmune disorders, indicating that similar genes are perturbed in related directions **(Figure 4B)**. In addition, we found that proteostasis proteins associated with neurodegenerative diseases are often perturbed in opposite directions to cancers and autoimmune diseases **(Figure 4B)**.

Further exploration of the functional implications of the proteostasis signatures reveal disease pathways modulated by proteostasis. In cancer, upregulated proteostasis proteins are enriched in cell cycle pathways **(Figure 4C)**, leading to aberrant activation of cell cycle proteins that promote cancer proliferation (25). Based on our analysis, the activation of cell cycle pathways may largely involve APC/C-dependent triggering of metaphase to anaphase transition during mitosis **(Figure 4C)**. This finding corroborates previous description of the pivotal role of APC/C in cell division and cell viability, which is implicated in tumorigenesis (26) – supporting the motivation for existing efforts seeking to target APC/C for therapeutic benefit(26) .

FOXO-mediated transcription, reported to be tumor suppressing (27, 28), was also found to be downregulated due to dysregulated proteostasis in cancers. Like cancers, cell cycle pathways have also been reported to be dysregulated in autoimmune diseases(29). This may likely be regulated by faulty disease-related proteostasis regulation of the pathway as shown in our results **(Figure 4C)**, and may suggest that the proteostasis dysregulation underlies the associated risk between the two disease types via cell cycle activation. Nevertheless, autoimmune diseases also present hyperactivation of the innate immune system **(Figure 4C)**, which breaks the inactivity of auto-reactive cells in the adaptive immune system, leading to autoimmune diseases (30).

The disease-induced changes in the proteostasis system affect several key cellular processes in neurodegenerative diseases. We found that the unfolded protein response (UPR) is activated, likely as a natural reaction to the accumulation of aggregation-prone and damaged proteins. Other dysregulated pathway include those involved in protein clearance, such as lysosome formation, autophagy, endocytosis, and recycling, alongside MITF-M-regulated pathways critical for maintaining brain function, which are often compromised in such conditions (31). Additionally, this disruption impacts ion channel transport and insulin signalling, further contributing to the disease pathology.

### Proteostasis perturbations in disease onset and progression

Next, we studied proteostasis perturbations on a temporal scale over the course of disease staging in 6 diseases (3 neurodegenerative diseases and 3 cancers) for which staging data are available: Alzheimer’s disease (AD), Parkinson’s disease (PD), Huntington’s disease (HTT), lung cancer, kidney cancer, and pancreatic cancer **(Figure 5)**. For each disease, differential gene expression analysis was carried out by disease stage against healthy controls. Our results reveal that proteostasis perturbations, regardless of upregulation or downregulation, occurr progressively in neurodegenerative diseases but early in cancers **(Figure 5)**. These trends were conserved across all diseases included in this analysis for both disease types.

**Figure 5:**
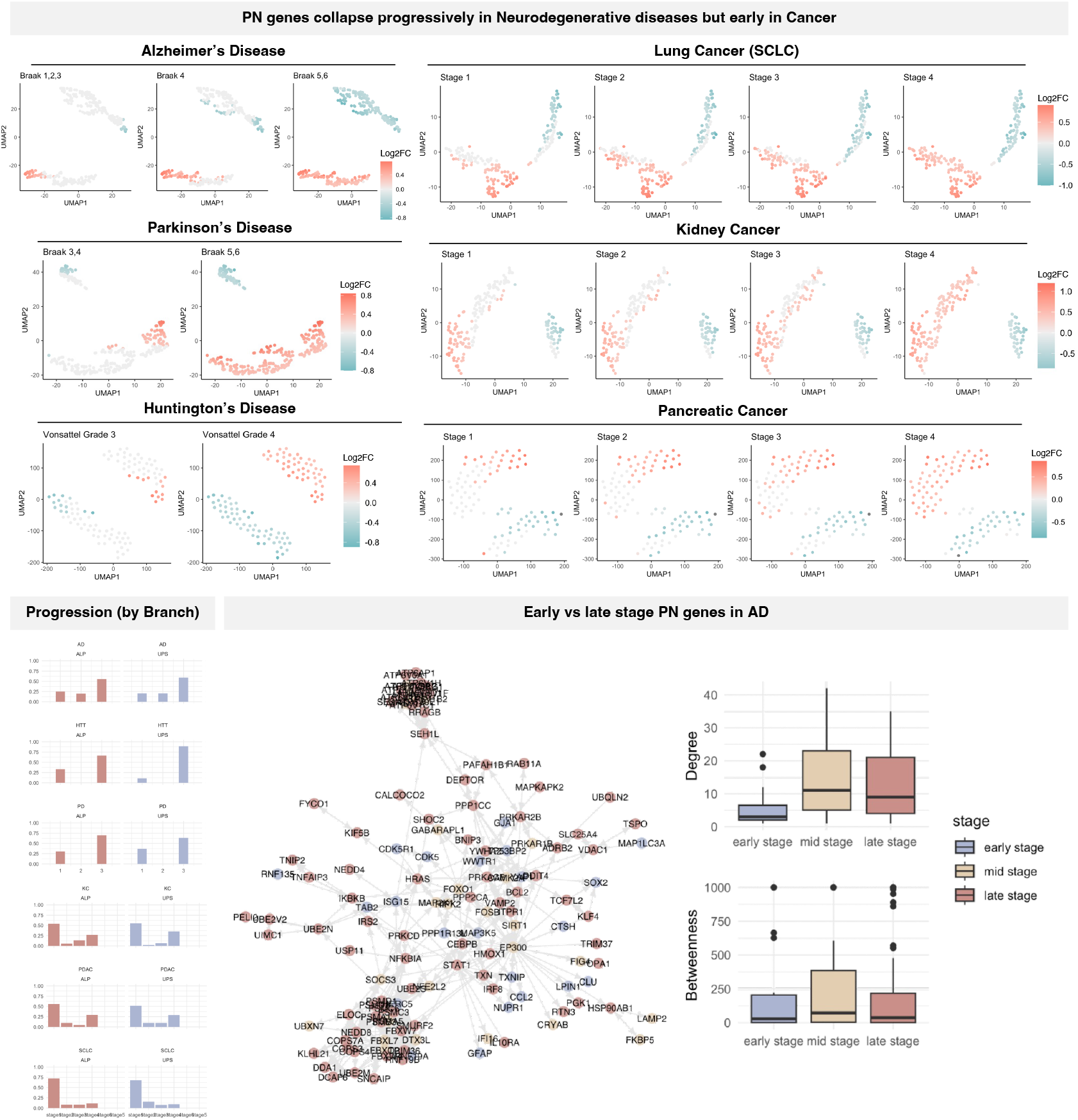
Proteostasis perturbation over disease onset and progression. Patient samples from each disease was compared against against healthy controls. Differential gene analysis reveals that perturbation of the proteostasis network (PN) occurred progressively in neurodegenerative diseases but early in cancers. Furthermore, despite ALP and UPS perturbations being indicative of disease states in both disease types, their occurrence happens at different stages of disease development. In AD, we find that early-stage AD proteostasis proteins are likely to be seeds that trigger a proteostasis collapse cascade in AD via affecting key regulatory proteins in mid-stages.

Further findings from this disease progression analysis revealed that although both ALP and UPS perturbations are indicative of disease states in both neurodegenerative diseases and cancer, they occur at different stages of disease progression. A larger proportion of affected ALP and UPS proteins are perturbed in early stages of the 3 cancers studied, but only at late stages for the 3 neurodegenerative diseases studied **(Figure 5)**. This result is in line with existing observations that damaged proteins are accumulated in neurodegenerative diseases over the course of aging that leads to toxicity, while cancer cells hijack the proteostasis system to enable survival and proliferation.

We further studied how proteostasis perturbations spread across disease progression. We hypothesized that the proteostasis proteins perturbed at earlier stages are central regulators of the proteostasis network, resulting in downstream disarray. To test this possibility, we mapped the perturbation of the proteostasis network in AD. We then quantified and compared the degree and betweenness centrality of the proteostasis proteins involved in each stage. The degree measures the number of connections a protein within the network, and betweenness measures the extent to which a protein lies on the shortest path between protein pairs within the network. Proteins with high degree and betweenness are likely to play key regulatory roles in their functional networks. Based on these metrics, we found that, unlike our initial hypothesis, proteostasis proteins perturbed in mid-stage AD (Braak 2/3) are most central in the AD proteostasis network **(Figure 5)**. This result prompts further investigations on whether early-stage AD proteostasis proteins could be seeds that affect regulatory proteins contributing to proteostasis collapse in later stages.

### Proteostasis perturbations due to smoking are associated with disease risk

We further investigated if smoking, a key risk factor for many chronic diseases, may increase disease risk via the proteostasis network. For this analysis, we carried out differential gene expression analysis between smokers and non-smokers with no reported diseases. We then compared the differentially expressed genes in smokers against 8 disease gene sets: chronic obstructive pulmonary disease (COPD), lung cancer, breast cancer, coronary heart disease, ulcerative colitis, endometrial cancer, endometriosis, and PD. It has been reported that smoking is a risk factor for the development of COPD (32, 33), lung cancer (34), breast cancer (35, 36), and coronary heart disease (37, 38). In contrast, smoking has been reported to reduce the risk for ulcerative colitis (39, 40), endometrial cancer (41-43), endometriosis (44-46), and PD(47-49). According to our hypothesis, we expected to find a higher similarity in perturbed proteostasis genes in smokers with diseases with increased risk due to smoking (hereafter referred to as ‘at-risk’ diseases), and a lower level of similarity

between smokers and diseases with lowered risk due to smoking (hereafter referred to as ‘reduced-risk’ diseases).

Our results show that proteostasis perturbations are more similar between smokers and patients with at-risk diseases and lower between smokers and patients with reduced-risk diseases **(Figure 6A)**. The Jaccard index was used to quantify similarity between perturbed proteostasis proteins due to smoking and proteostasis proteins within each disease gene set **(Materials and Methods)**, then normalized for plotting in **Figure 6A**. We found that quantifying similarities between smoking-impacted proteostasis was more indicative of risk of disease than smoking-impacted kinases or transcription factors **(Figure 6A)**, both of which are protein groups strongly associated with disease (as discussed earlier).

**Figure 6:**
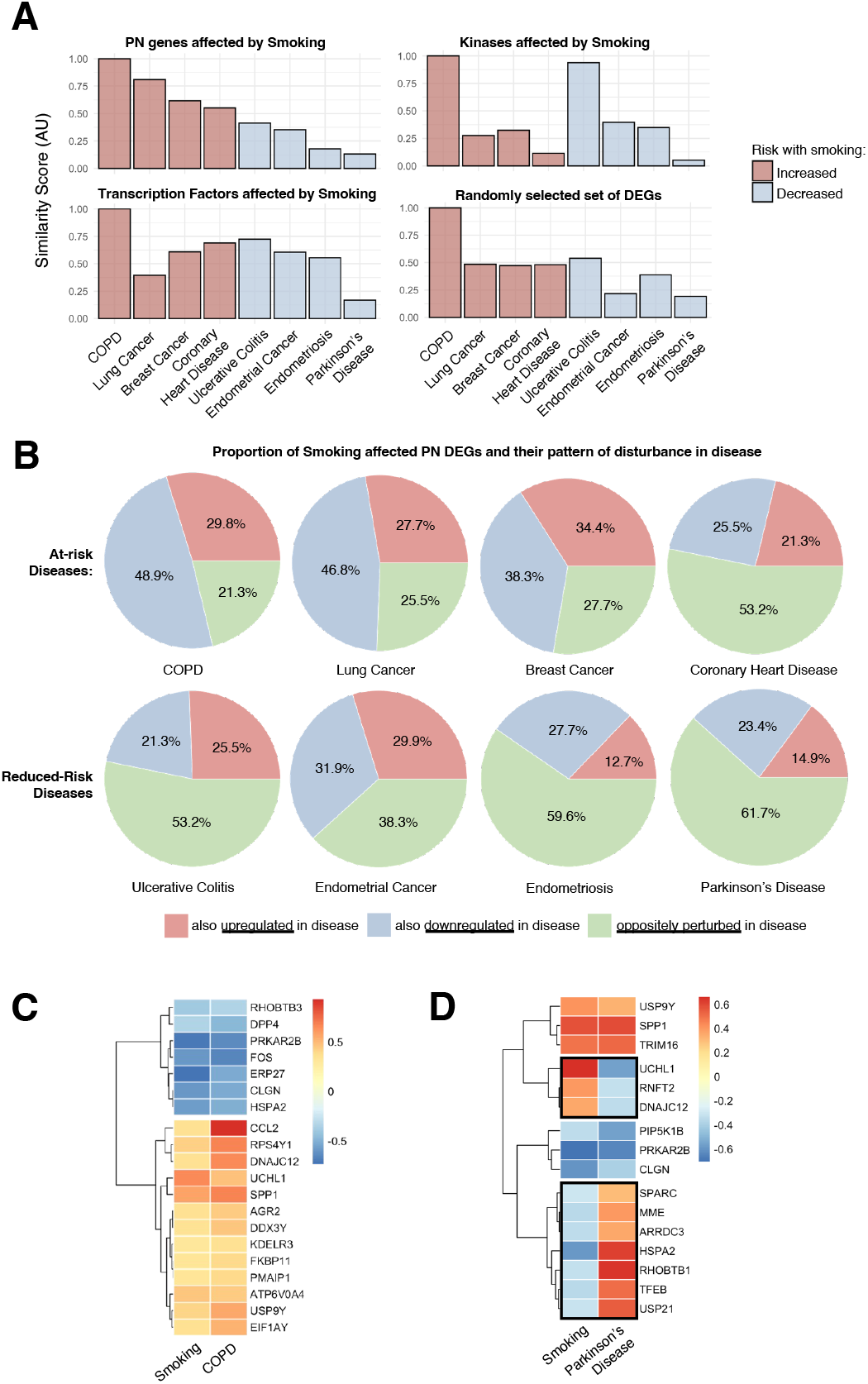
Proteostasis perturbations due to smoking are indicative of disease risk. **(A)** Smokers present a higher similarity of proteostasis perturbations with at-risk diseases compared to reduced-risk diseases. Computing similarities of proteostasis proteins is more indicative of disease risk as compared to smoking-impacted kinases, transcription factors, or a random sample of differentially expressed genes. **(B)** At-risk diseases have a higher directional similarity of their perturbed proteostasis proteins with smoking. In contrast, reduced-risk diseases have a large proportion of perturbed proteostasis proteins that are deregulated in the opposite direction. **(C)** Genes encoding proteostasis proteins are perturbed similarly in smokers and patients with COPD. Genes similarly perturbed between smoking and COPD are likely to be contributive toward increasing COPD risk and onset. **(D)** Proteostasis proteins corresponding to genes perturbed in smokers and patients with PD. Proteostasis proteins oppositely perturbed between smoking and PD are likely to be protective against PD.

We then compared the pairwise directional similarity of proteostasis perturbation between the overlapping smoking-perturbed proteostasis and disease-associated proteostasis perturbations. Our comparison revealed that smoking results in a larger directional similarity (i.e. proteostasis genes upregulated due to smoking, are also upregulated in disease; or proteostasis genes downregulated due to smoking, are also downregulated in disease) with at-risk diseases vis-à-vis reduced-risk diseases that have a large proportion of their proteostasis being perturbed in opposite directions **(Figure 6B)**.

These results suggest that the proteostasis similarly perturbed between smoking and at-risk diseases are likely to contribute toward increasing disease risk and the development of disease pathologies. Given this observation, we further investigated the proteostasis proteins perturbed similarly in smokers and patients with COPD (at-risk disease with highest similarity). For example, CCL2, an extensively studied protein target in COPD (50, 51), is upregulated due to smoking **(Figure 6C)**. Similarly, proteostasis oppositely perturbed between smoking and reduced-risk diseases are likely to be protective against disease development. Hence, we investigated PD (reduced-risk disease with largest dissimilarity) in more detail. *UCHL1* is an example of a very highly over-expressed gene in smokers, which contrasts the downregulation of *UCHL1* observed in PD patients **(Figure 6D)**. It has been reported to be a susceptibility gene for PD and proposed as a potential target for therapy (52), whose downregulation contributes to protein aggregation in Lewy bodies (53) – a hallmark of PD pathology. Based on this, it is likely that the upregulation of *UCHL1* due to smoking protects against *UCHL1* loss-of-function that predisposes the cells to PD related symptoms. Moving forward, it will be interesting to further explore the other genes **(Figure 6C,D)** that present this trend to uncover key contributors to disease vulnerability thus supporting efforts in preventive care.

## Discussion

In this work, we analysed the association of the proteostasis network with disease, and identified the corresponding proteostasis signatures. We also identified temporal patterns of proteostasis involvement during disease development. We further uncovered how risk factors associated with lifestyle, such as smoking, greatly impact the proteostasis network, likely priming for disease cells with high vulnerability.

Profiling proteostasis signatures has the potential to increase our understanding of the proteostasis mechanisms driving disease aetiologies. We note, however, that caution must be exercised when interpreting these proteostasis signatures, as association does not necessarily imply causation, and extensive effort needs to go into validating these mechanisms so as to develop robust and effective interventions.

Overall, we anticipate that approaches based on the analysis of proteostasis signatures could lead to identifying therapeutic avenues by preventing proteostasis collapse, or restoring perturbed proteostasis functions.

### Materials and Methods Proteostasis network

A comprehensive list of proteostasis network components (13, 14) were obtained from the Proteostasis Consortium (https://www.proteostasisconsortium.com/).

### Disease gene sets

The top 500 genes for each disease, based on PandaOmics (54), were determined to be disease associated and made up the gene set. This ranking was derived by comparing disease samples to tissue-matched controls in PandaOmics v2.0. The final scores used for ranking are a result of aggregating multiple omics inputs for each gene using a neural network. The multi-omics input include: (i) mRNA expression (level of differential gene expression in disease versus control), (ii) interactome community (the density of known targets, disease-related genes, and differentially expressed genes in its protein-protein interaction network), (iii) causal inference (estimating the number of genes regulated by similar transcription factors), (iv) overexpression (characterizing the effects of gene knock-in/knock-out on cell lines), (v) mutated disease sub-modules (assessing gene relevance based on OMIM, ClinVar, and Open Targets data), (vi) mutations (a combined score from genome and transcriptome-wide association studies), (vii) pathway analysis (evaluating a gene’s role in Reactome pathways using iPANDA and transcriptomic data), (viii) network neighbors (based on the number of directly connected differentially expressed genes in the protein interaction network), and (ix) disease relevance (aggregated scores from OMIM, ClinVar, and Open Targets). The proteostasis subset of each disease gene set was obtained by finding the intersection of our full list of proteostasis components with each disease gene set. All disease gene sets are available in **Table S4**.

### Over-representation analysis of proteostasis proteins

A list of kinases, transcription factors, and ion channels **(Table S5)** were obtained from the PandaOmics database that annotated more than 20,000 proteins. Over-representation analysis was carried out using the hypergeometric test that measures the statistical significance of each group of proteins being over-represented in the disease gene set. P-values were plotted on a -log(p-value) scale, with higher values representing stronger significance.

### Over-representation analysis of proteostasis network branches and classes

The hypergeometric test was used to quantify the enrichment of every branch/class within the disease gene set for each diseases. A cutoff of p-value <0.01 was used in the profiling presented in **Figure 5**.

### Unsupervised clustering of diseases

The Jaccard Index was used to calculate similarity scores between diseases. This similarity matrix was used for hierarchical clustering (hclust function in R) which generated 4 clusters **(Figure 6A)**.

### Pathway enrichment analysis

Pathway enrichment analysis was carried out by calculating the probability of a disease gene set being over-represented in a pathway. The Reactome pathways were used for this analysis. A cut-off of false discovery rates (FDR) < 0.05 and more than 2 genes per pathway were applied. For plotting, the most upregulated and downregulated (up to 10 each) were included.

### Disease staging transcriptomic datasets

Only diseases for which transcriptomic datasets with disease staging information that was readily available were included in this study. The datasets used were: GSE48350 and GSE84422 (AD); GSE49036 and GSE42966 (PD); GSE64810 and GSE79666 (HTT); GSE30219 (lung cancer); GSE53757, GSE76207, and GSE126964 (kidney cancer); GSE62452 (pancreatic cancer).

### Smoking transcriptomic datasets

Datasets that included samples from healthy controls that were smokers and non-smokers were obtained to study the impact of smoking on the proteostasis network. Datasets obtained and used include GSE22047 and GSE108134.

### Differential expression analysis

Differential expression analysis was used to quantify expression differences between conditions: disease stage vs healthy control, or smokers vs non-smokers, for disease staging and smoking transcriptomic datasets respectively. Each analysis done using the Limma method (55).

### Similarity scores

The Jaccard index was used to calculate similarity scores. It measures the ratio of the overlap between the perturbed proteostasis genes due to smoking and the proteostasis genes within each disease gene set to their union.

## Supplementary Material

**Table S1.** List of diseases and their corresponding disease groups.

| Disease Group | Disease Name |
| --- | --- |
| autoimmune | Celiac Disease |
| autoimmune | Crohns Disease |
| autoimmune | Lupus |
| autoimmune | Rheumatoid Arthritis |
| autoimmune | Ulcerative Colitis |
| cancer | Breast Cancer (HER2+) |
| cancer | Breast Cancer (TN) |
| cancer | Cervical Cancer |
| cancer | Colorectal Cancer |
| cancer | Endometrial Cancer |
| cancer | Kidney Cancer |
| cancer | Liver Cancer (HCC) |
| cancer | Lung Cancer (NSCLC) |
| cancer | Lung Cancer (SCLC) |
| cancer | Melanoma |
| cancer | Ovarian Cancer |
| cancer | Pancreatic Cancer |
| cardiovascular | Coronary Heart Disease |
| cardiovascular | Dilated Cardiomyopathy |
| cardiovascular | Hypertrophic Cardiomyopathy |
| cardiovascular | Ischemic Cardiomyopathy |
| endocrine | Type I Diabetes |
| endocrine | Type II Diabetes |
| neurodegenerative | Alzheimers Disease |
| neurodegenerative | CJD |
| neurodegenerative | Huntingtons Disease |
| neurodegenerative | Parkinsons Disease |
| neurodegenerative | Picks Disease |
| reproductive | Endometriosis |
| reproductive | PCOS |
| respiratory | Asthma |
| respiratory | COPD |

**Table S2.** Proteostasis network branches.

| Branch Code | Proteostasis Branch Name |
| --- | --- |
| A | Autophagy |
| U | UPS |
| Pn | PN Regulation |
| Ex | Extracellular Proteostasis |
| Ct | Cytonuclear Proteostasis |
| Nu | Nuclear Proteostasis |
| Mt | Mitochondrial Proteostasis |
| Er | ER Proteostasis |

**Table S3.** Proteostasis network classes.

| **Class Code** | **Proteostasis Class Name** |
| --- | --- |
| Ge | Autophagy Gene Exp |
| Ca | Chaperone-Mediated Autophagy |
| Uk | Functions Unknown |
| Pa | Proteosome associated |
| Si | Pre-initiation Signalling |
| Ie | Autophagophore Initiation and elongation |
| Ss | Substrate selection |
| Lf | Autophagosome closure & Lysosome fusion |
| Lc | Lysosome catabolism |
| Lr | Lysosome reformation |
| Ub | Ubiquitin binding |
| Up | Ubiquitin proteins |
| E1 | E1 ligases |
| E3 | E3 ligases |
| E2 | E2 ligases |
| Du | DUBS and UBL demodifiers |
| Tf | Transcription Factor |
| Re | TF regulator |
| C | Chaperone |
| P | Protease |
| Tr | Translation regulator |
| M | Substrate Maturation |

**Table S4**. Disease gene sets for all 32 diseases studied (see separate Excel file).

**Table S5**. List of kinases, transcription factors, and ion channels used as controls (see separate Excel file).

